# Biased agonists of the chemokine receptor CXCR3 differentially drive formation of G_αi_:β-arrestin complexes

**DOI:** 10.1101/2020.06.11.146605

**Authors:** Kevin Zheng, Jeffrey S. Smith, Anmol Warman, Issac Choi, Jaimee N. Gundry, Thomas F. Pack, Asuka Inoue, Marc G. Caron, Sudarshan Rajagopal

## Abstract

G-protein-coupled receptors (GPCRs), the largest family of cell surface receptors, signal through the proximal effectors G proteins and β-arrestins to influence nearly every biological process. Classically, the G protein and β-arrestin signaling pathways have largely been considered separable. Recently, direct interactions between G_α_ protein and β-arrestin have been described and suggest a distinct GPCR signaling pathway. Within these newly described G_α_:β-arrestin complexes, G_αi/o_, but not other G_α_ protein subtypes, have been appreciated to directly interact with β-arrestin, regardless of canonical GPCR G_α_ protein subtype coupling. However it is unclear how biased agonists differentially regulate this newly described G_αi_:β-arrestin interaction, if at all. Here we report that endogenous ligands (chemokines) of the GPCR CXCR3, CXCL9, CXCL10, and CXCL11, along with two small molecule biased CXCR3 agonists, differentially promote the formation of G_αi_:β-arrestin complexes. The ability of CXCR3 agonists to form G_αi_:β-arrestin complexes does not correlate well with either G protein signaling or β-arrestin recruitment. Conformational biosensors demonstrate that ligands that promoted G_αi_:β-arrestin complex formation generated similar β-arrestin conformations. We find these G_αi_:β-arrestin complexes can associate with CXCR3, but not with ERK. These findings further support that G_αi_:β-arrestin complex formation is a distinct GPCR signaling pathway and enhance our understanding of biased agonism.

## Introduction

GPCRs are receptors that enable cells to sense and respond appropriately to hormonal and environmental signals. Targeting GPCR signaling has proven therapeutic benefit as approximately 30% of all FDA approved medications target GPCRs (*1*). Classically, each GPCR couples to distinct G_α_ protein families, such as G_αs_, G_αi_, G_αq_ or G_α12/13_, as well as β-arrestins (*2, 3*). These transducer proteins are utilized by nearly every GPCR to translate and integrate extracellular stimuli into intracellular signals. It has been largely considered that G proteins and β-arrestins comprise separable GPCR signaling pathways, with β-arrestins acting both as independent signaling scaffolds for downstream effectors while also serving as critical negative regulators of G protein signaling (*4-8*). In addition, recent evidence suggests that there is direct communication between G_α_ proteins and β-arrestins, specifically G_αi_ and β-arrestins, that can regulate cellular function such as ERK signaling. It is known that GPCRs can form “megaplex” signaling complexes, in which a GPCR is able to bind to both G protein and β-arrestin (*9, 10*). We recently reported that agonist treatment of GPCRs, regardless of canonical G_α_ protein:GPCR coupling, can catalyze the formation of G_αi_:β-arrestin signaling complexes (Smith and Pack et al., submitted). G_αi_ appears to play a unique role in influencing β-arrestin signaling relative to other G protein isoforms. Indeed, such G_αi_:β-arrestin complexes demonstrated the ability to form G_αi_:β-arrestin:GPCR complexes with the V_2_R and β_2_AR. These signaling complexes potentially have widespread physiological and therapeutic implications. However, it remains unclear if different agonists for the same GPCR form G_αi_:β-arrestin complexes in a conserved fashion, if G_αi_:β-arrestin complex formation strongly correlates with other established arms of GPCR activation, or if Gαi:β-arrestin complexes interact with receptors or signaling effectors such as ERK in a conserved manner.

It is currently unknown if formation of G_αi_:β-arrestin complexes closely correlate with canonical GPCR activation events, such as G protein recruitment, G protein signaling (e.g., cAMP inhibition), or β-arrestin recruitment. Findings that some agonists can bind the same receptor and differentially activate downstream distinct signaling pathways, a well-established phenomenon (*11*) referred to as biased agonism, has typically shown that G proteins and β-arrestins promote signaling through relatively independent pathways (*12, 13*). Both synthetic G protein- and β-arrestin-biased agonists have been identified for many GPCRs, although only a few GPCRs have been shown to have multiple endogenous ligands that act as biased agonists. One of the best examples of endogenous biased agonism is in the chemokine system, where the majority of chemokine receptors bind more than one chemokine, and many chemokines act as biased agonists of their receptors (*14, 15*). We have previously studied the receptor CXCR3, which has three endogenous ligands, CXCL9, CXCL10, and CXCL11, and is involved in many diseases, including cancer, cutaneous infection, atherosclerosis, hypersensitivity reactions, and autoimmune disorders (*16-20*). We previously demonstrated that CXCL11 is β-arrestin-biased relative to CXCL9 and CXCL10 (*15, 21*). In addition, CXCR3 has known small molecule biased agonists, such as VUF10661 and VUF11418 (*22, 23*), that we have previously validated as β-arrestin- and G protein-biased agonists, respectively (*24*).

To determine the potential linkage between canonical and noncanonical GPCR signaling we studied three well-established CXCR3 signaling pathways: cAMP inhibition (G_αi_ protein signaling), G protein recruitment to CXCR3, and β-arrestin recruitment to CXCR3. We also tested the ability of these five agonists to form G_αi_:β-arrestin complexes. We found that these agonists had distinct G protein signaling, β-arrestin recruitment, and G protein recruitment profiles. However, none of these profiles clearly corresponded to the ability to form G_ai_:β-arrestin complexes, consistent with this complex being distinct from canonical pathways. Using a panel of β-arrestin biosensors, we found that ligands that promoted G_ai_:β-arrestin complexes generated similar β-arrestin conformations to one another. This study provides critical insights into GPCR activation events that govern G_αi_:β-arrestin complex formation and expands our concept of biased agonism.

## Results

### Assessment of canonical CXCR3 signaling

We first compared the activity of five CXCR3 agonists, CXCL9, CXCL10, CXCL11, VUF10661, and VUF11418 in multiple assays: G_αi/o_ protein recruitment, G protein signaling (inhibition of cAMP), and β-arrestin recruitment. Importantly, G protein *recruitment* and G protein *signaling* (such as cAMP inhibition in the case of G_αi_) may not correlate with one another due to differences in G protein pre-coupling or other mechanisms (*25, 26*). Furthermore, as CXCR3 is a canonically G_αi_-coupled receptor, either G protein recruitment or signaling may correlate better with G_αi_:β-arrestin complex formation. First, we tested the ability of the panel of CXCR3 agonists to recruit G_αi_ subunits to CXCR3 using a previously described nanoBiT system (Fig. 1a) (*26, 27*). G_αi_ family members G_αi-1_ and G_αo_, but not G_αs_, were recruited to CXCR3 following treatment with CXCL11, the β-arrestin-biased agonist VUF10661, and the G protein-biased agonist VUF11418 (Fig. 1b,c, Supp. Fig. 1a,b). CXCL9 or CXCL10 treatment did not induce G_αi/o_ protein recruitment to CXCR3 as robustly as did CXCL11, VUF10661, and VUF11418 (Fig. 1b,c, Supp. Fig. 1a). The observed interaction of G_αi_ with CXCR3 was sensitive to pertussis toxin pretreatment (Supp. Fig. 1c,d), which promotes ADP ribosylation of cysteine 352 in helix 5 of G_αi_, and significantly attenuated the observed interaction between CXCR3 and G_αi/o_. Overexpression of β-arrestin, which sterically hinders G proteins from interacting with GPCRs, also significantly attenuated G protein interaction with CXCR3 (Supp. Fig. 2a). Combining pertussis toxin pretreatment and overexpression of β-arrestin completely inhibited G protein recruitment to CXCR3 (Supp. Fig. 2b). Neither overexpression of β-arrestin, pretreatment with pertussis toxin, nor a combination of the two significantly altered CXCR3’s inability to recruit G_αs_ (Supp. Fig. 2c,d).

**Figure 1:**
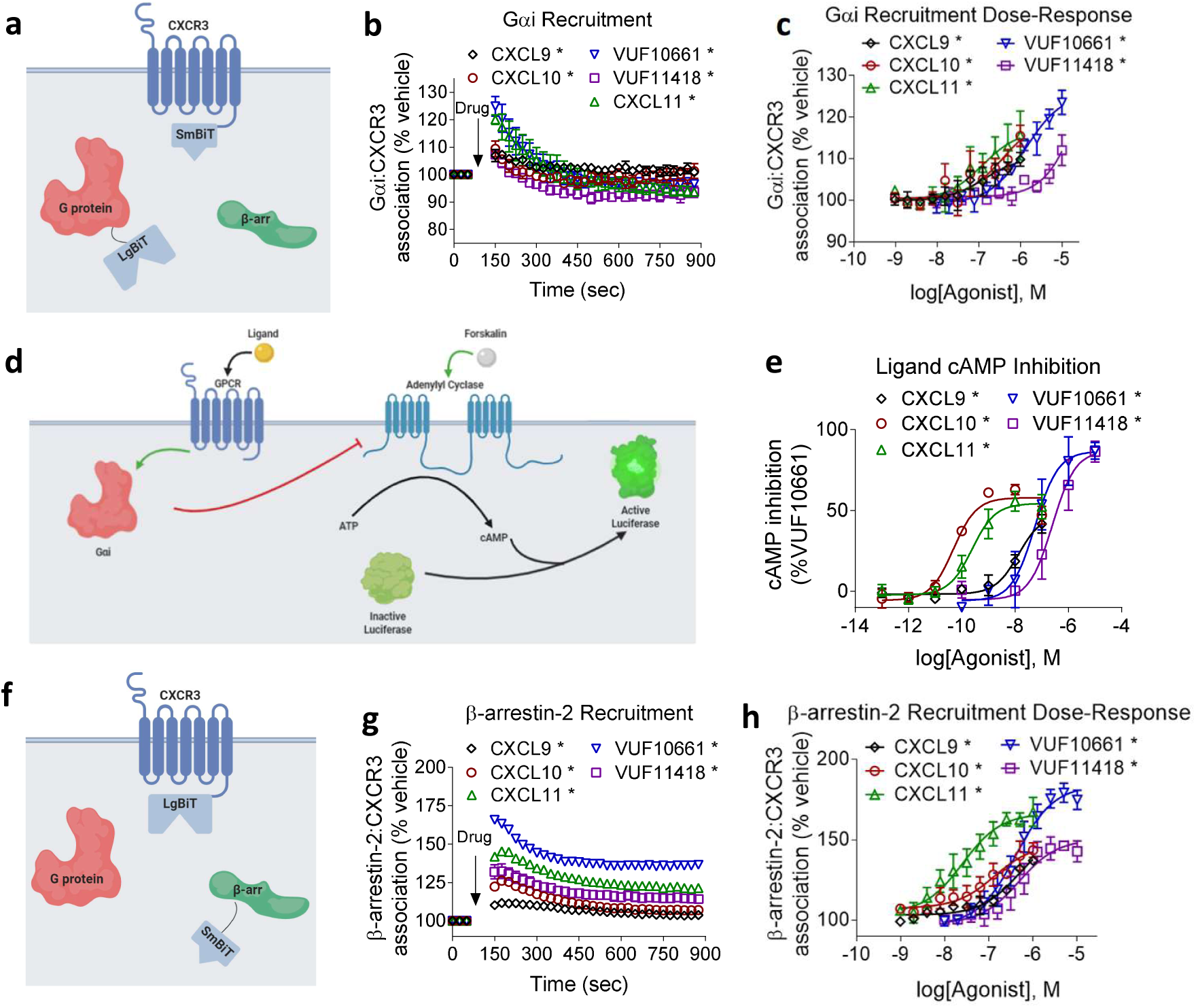
G_αi_ protein recruitment, cAMP inhibition, and β-arrestin-2 recruitment by CXCR3 biased agonists. A) Arrangement of nanoBiT luciferase fragments to assess G protein recruitment. B) HEK 293T transiently expressing CXCR3-smBiT and G_αi_-LgBiT were treated with either CXCL9 (100nM), CXCL10 (100nM), CXCL11 (100nM), VUF10661 (1μM), VUF11418 (1μM), or vehicle at the specified concentration and analyzed for 750 seconds after an initial pre-read. C) G_αi_ recruitment to CXCR3 following agonist treatment at indicated concentrations one-minute post ligand treatment. D) Schematic illustrating G_αi_-regulated cAMP inhibition assay. Prior to treatment with CXCR3 biased agonists, cellular cAMP was increased with 10μM of forskolin. E) HEK 293T transiently expressing cAMP-activated modified firefly luciferase and CXCR3 were treated with vehicle or the indicated concentrations of CXCL9 (100nM), CXCL10 (100nM), CXCL11 (100nM), VUF10661 (1μM), or VUF11418 (1μM). cAMP inhibition is displayed as change relative to VUF10661. F) Arrangement of luciferase fragments CXCR3-LgBiT and smBiT-β-arrestin-2 to assess β-arrestin-2 recruitment to CXCR3. G) HEK 293T transiently expressing CXCR3-LgBiT and smBiT-β-arrestin-2 were treated with either CXCL9 (100nM), CXCL10 (100nM), CXCL11 (100nM), VUF10661 (1μM), VUF11418 (1μM), or vehicle and analyzed for 750 seconds after an initial pre-read. H) Assessment of β-arrestin-2 recruitment following agonist treatment at the indicated concentrations six-minutes post ligand treatment. *P<0.05 by two-way ANOVA, Dunnett’s post hoc analysis with a significant difference relative to pretreatment. n = 3-4, graphs show mean ± s.e.m.

Next, we tested the ability of CXCR3 to promote canonical G_αi_ signaling, which we monitored as a reduction of intracellular cAMP with the Glosensor™ (Promega) cAMP luminescent biosensor (Fig. 1d). Consistent with the predicted signaling properties of G_αi_-coupled CXCR3, all five ligands reduced intracellular cAMP in a dose-dependent fashion, with potencies that mirrored the affinities of each agonist for CXCR3. The two small-molecule agonists appeared to have higher efficacy than the endogenous chemokines (Fig. 1e). This reduction in cAMP was promoted by CXCR3, as cells transiently transfected with empty vector did not display the same reduction when treated with the agonist panel (Supp. Fig. 3a-e). Interestingly, despite the inability to robustly recruit G_αi/o_ by CXCL9 and CXCL10, both chemokines were able to inhibit cAMP to similar degrees relative to the other three agonists. These data indicate that G protein recruitment and G protein signaling as regulated by CXCR3 are discernable properties.

We then assessed CXCR3 β-arrestin recruitment induced by the agonist panel using a nanoLuc complementation assay (Fig 1f). Consistent with prior observations, all five agonists were able to recruit β-arrestin. VUF10661 had the highest efficacy for β-arrestin recruitment followed by CXCL11, VUF11418, CXCL9, and CXCL10 (Fig. 1g). Importantly, VUF10661 and CXCL11 promoted significantly more β-arrestin recruitment than the other three agonists as assessed by dose-response (Fig. 1h). Pertussis toxin pretreatment did not affect agonist-induced β-arrestin recruitment to CXCR3, consistent with previous observations at other GPCRs (Smith and Pack et al., submitted) (Supp. Fig 4a-e).

### Biased agonists of CXCR3 differentially promote formation of G_αi_:β-arrestin complexes

We then examined whether these five CXCR3 agonists could promote the formation of G_αi_:β-arrestin complexes. We previously demonstrated that an association between G_αi_ family proteins and β-arrestins could be catalyzed by ligand treatment of a variety of GPCRs, even those that do not classically couple to G_αi_ such as the vasopressin receptor (V_2_R), β_2_-adrenergic receptor (β_2_AR), and neurotensin receptor type 1 (NT_1_R) (Smith and Pack et al., submitted). However, it is unknown if biased agonists would differentially form G_αi_:β-arrestin complexes. To address this, we treated HEK293 cells overexpressing CXCR3, G protein-LgBiT, and SmBiT-β-arrestin with our agonist panel and tested for G_αi_ and β-arrestin association (Fig. 2a). Treatment of CXCR3 with CXCL11, the β-arrestin-biased agonist VUF10661, or the G protein-biased agonist VUF11418 resulted in association of G_αi_ and β-arrestin (Fig. 2b,c). Similarly, treatment of CXCR3 with CXCL11 and VUF10661 resulted in G_αo_ and β-arrestin association (Supp. Fig. 5a). Notably, neither CXCL9 nor CXCL10 treatment of CXCR3 resulted in appreciable G_αi_ or G_αo_ association with β-arrestin (Fig. 2b, Supp. Fig. 5a). Consistent with our previously reported results with other receptors, none of the agonist treatments promoted association of β-arrestin with the G_αs_, G_αq_, G_α12_ proteins (Supp. Fig. 5b-d) (Smith and Pack et al., submitted). As many of these chemokines have alternative receptors, we also tested the effects of these ligands on cells transfected solely with the β-arrestin and G_αi_ components and not CXCR3. We did not observe a significant increase for any ligand aside from CXCL9 (Supp. Fig. 5e), which is likely through its activity on an endogenously expressed receptor. This is consistent with the ability of CXCL9 to reduce cAMP in non-CXCR3 transfected cells (Supp. Fig. 3a). Consistent with our previously reported observations, pertussis toxin pretreatment nearly eliminated G_αi_ and β-arrestin association at all three agonists that promoted complex formation (CXCL11, VUF10661, and VUF11418) (Fig. 2d-e). Compared with G_αi_ and β-arrestin recruitment, pertussis toxin pretreatment was only able to attenuate G_αi_:β-arrestin formation and G_αi_ recruitment, but not β-arrestin recruitment.

**Figure 2:**
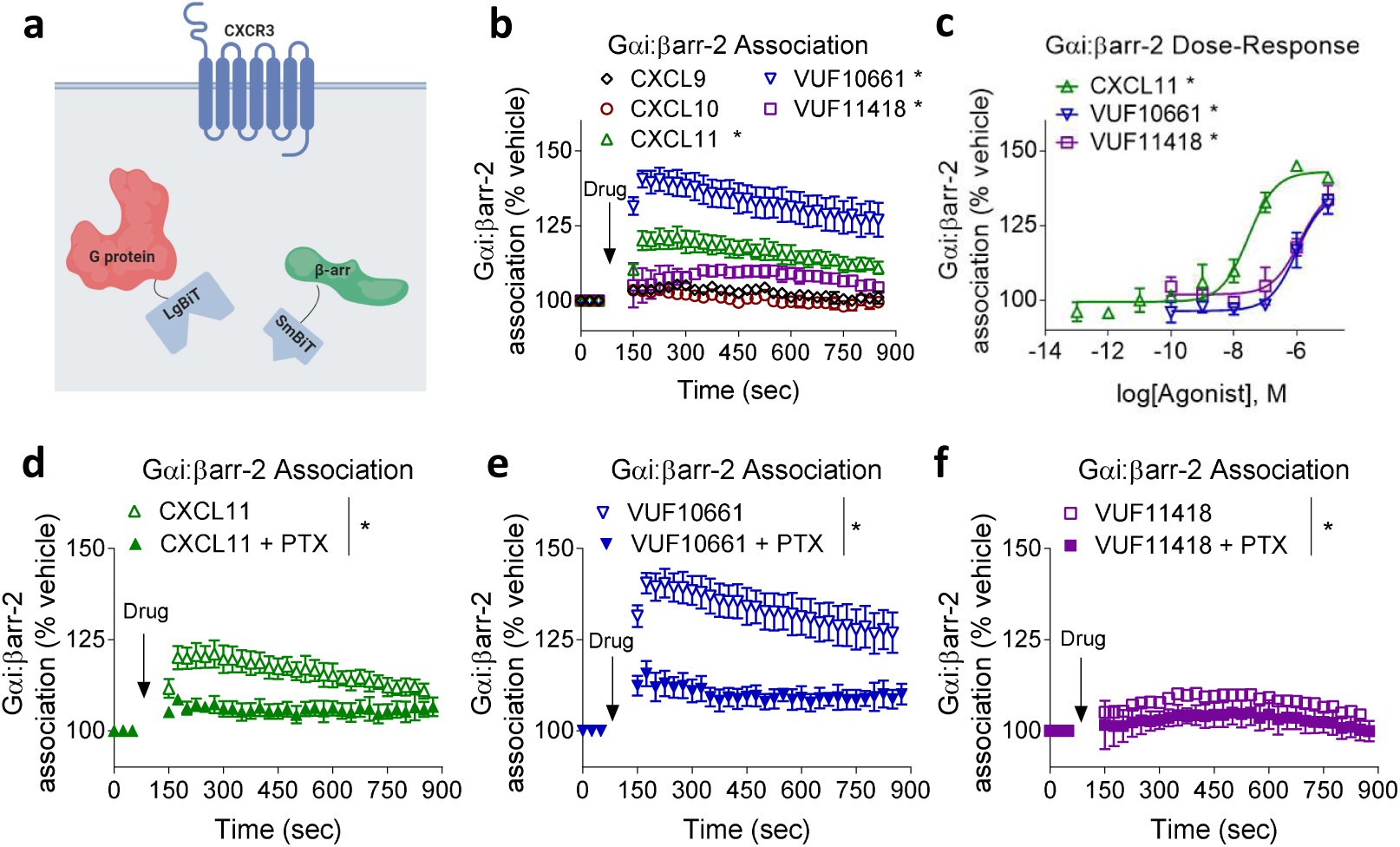
Biased agonists of CXCR3 differentially drive formation of G_αi_:β-arrestin complexes. A) Schematic of assay used to assess G_αi_:β-arrestin complex formation. B) HEK 293T cells transiently expressing CXCR3, smBiT-β-arrestin-2, and G_αi_-LgBiT were treated with either CXCL9 (100nM), CXCL10 (100nM), CXCL11 (100nM), VUF10661 (1μM), VUF11418 (1μM), or vehicle. G_αi_:β-arrestin complexes were observed with CXCL11, VUF10661, and VUF11418, but not with CXCL9 or CXCL10. C) Transfected HEK 293T cells expressing CXCR3, smBiT-β-arrestin-2, and G_αi_-LgBiT were treated with either CXCL11, VUF10661, VUF11418, or vehicle at the specified concentrations six-minutes post ligand treatment. HEK 293T cells transiently expressing CXCR3, smBiT-β-arrestin-2, and G_αi_-LgBiT were pretreated with pertussis toxin (200 ng/mL) and subsequently treated with either D) CXCL11, E) VUF10661, or F) VUF11418. In all cases, pertussis pretreatment resulted in significant attenuation of G_αi_:β-arrestin association relative to non-pertussis pretreated cells. *P<0.05 by two-way ANOVA, Dunnett’s post hoc analysis with a significant difference between treatments. n = 3-5, graphs show mean ± s.e.m.

### Assessment of CXCR3 biased signaling through G proteins, β-arrestins and G_αi_:β-arrestin complexes

We first qualitatively compared G_αi_:β-arrestin complex formation, β-arrestin recruitment, G protein recruitment and cAMP inhibition by constructing bias plots as previously described (*28, 29*) (Fig 3a-f). Bias plots allow for an assessment of assay amplification effects that can confound efforts to identify biased ligands (*7*). For example, because of second messenger amplification, a cAMP inhibition assay often demonstrates higher potencies and efficacies relative to a G protein recruitment assay. Such amplification can result in quantitative false positives, but can be easily visualized in bias plots. If a ligand does not appear to be biased using a bias plot, it is unlikely to be biased, as calculated bias factors can frequently have large errors (*28*). This is primarily because such bias plots are not prone to errors introduced from different fitting approaches. From such bias plots, we qualitatively observed that while the relationship between β-arrestin-2 recruitment and G_αi_:β-arrestin complex formation was relatively similar between CXCL11, VUF10661 and VUF11418, there was considerably more variance in the relationship between G_αi_ recruitment and G_αi_:β-arrestin complex formation.

**Figure 3:**
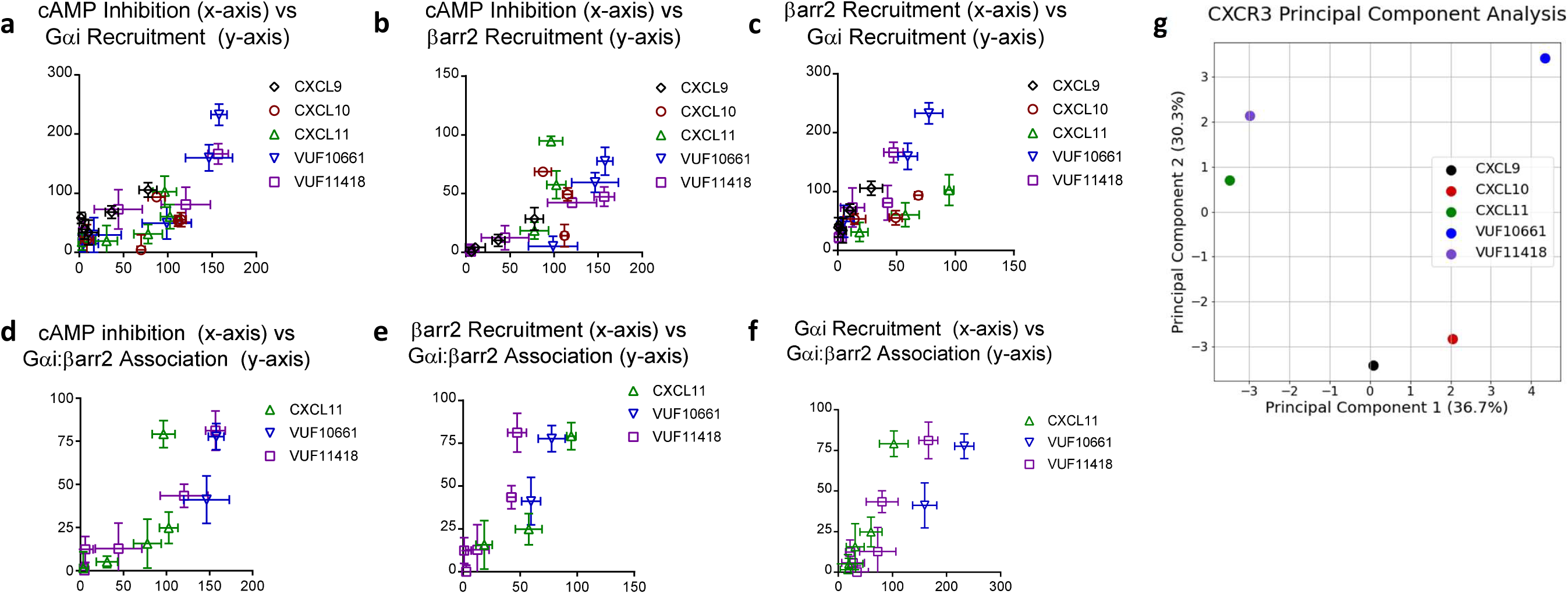
Bias plots for identification of biased ligands at CXCR3 in G protein recruitment, G protein signaling, β-arrestin-2 recruitment, and G_αi_:β-arrestin-2 association. As CXCL11 is the only full endogenous agonist for both G protein and β-arrestin pathways, we utilized it as the reference ligand. A) An equimolar comparison between cAMP inhibition and G_αi_ recruitment demonstrates a strong positive correlation without significant bias. B) An equimolar comparison between cAMP inhibition and β-arrestin-2 recruitment assay demonstrates a cAMP inhibition bias in VUF10661 and VUF11418. C) An equimolar comparison between β-arrestin-2 recruitment and G_αi_ recruitment demonstrates a G_αi_ recruitment bias in VUF10661 and VUF 11418. D) An equimolar comparison between cAMP and G_αi_:β-arrestin-2 association demonstrates a cAMP inhibition bias in VUF10661 and VUF11418. E) An equimolar comparison between β-arrestin-2 recruitment and G_αi_:β-arrestin association demonstrates a G_αi_:β-arrestin association bias in VUF 10661 and VUF 11418. F) An equimolar comparison between G_αi_ recruitment and G_αi_:β-arrestin association demonstrates a G_αi_ recruitment bias in VUF10661 and VUF11418. G) Principal Component Analysis (PCA) was used to assess CXCR3 agonist similarity based on signaling assays; computed principal components are visualized with the top two principal components. Principal Component 1 contributes to 36.71% of observed variation and Principal Component 2 contributes to 30.32% of observed variation. Points denote the composite response of a single ligand at varying doses.

To quantify these relationships further, we then calculated fitted curve parameters (EC_50_ and E_max_) based on the dose-response data utilized for the bias plots (Table 1). These parameters were then utilized to calculate the logarithm of ratios of relative intrinsic activities for each pair of pathways to determine bias factor values (Table 2). Consistent with the bias plots, the relationships between G_αi_:β-arrestin complex formation and β-arrestin recruitment show smaller bias factors than those of G_αi_ recruitment or cAMP inhibition. It should be noted that while both CXCL9 and CXCL10 promoted β-arrestin recruitment (Fig. 1g), neither promoted G_αi_:β-arrestin complexes. We subsequently performed linear regression of each ligand per bias plot to calculate an R^2 value, describing the linear similarity between two signaling pathways per agonist (Supp. Table 1). We then performed a principal component analysis (PCA), with the activity of each ligand at each pathway serving as input components. Our analysis allows for a 2-dimensional reduction of each ligand’s assessed signaling characteristics to evaluate agonist similarity. Utilizing this strategy, we visualized that the signaling of these five agonists fell into three profiles: one centered around CXCL9 and CXCL10, another around CXCL11 and VUF11418, and the last around VUF10661. Overall, this analysis demonstrates that these ligands display significant bias between their signaling pathways, suggesting that both ligands and pathways, including the recently appreciated G_αi_:β-arrestin pathway, are functionally distinct.

**Table 1:** Canonical and noncanonical signaling via CXCR3. Summary of fitting parameters. EC_50_ and E_max_ (expressed as % of CXCL11 signal) measurements for agonists calculated from a 3-parameter fit (y=Min + (Max-min)/(1+10^(LogEC_50_-X))). For G_αi_:β-arrestin complex formation, values were only calculated with CXCL11, VUF10661 and VUF11418, as the other agonists did not form a complex. If the 3-parameter fit of ligand-receptor interaction produced a poor fit, then the data was omitted from the table.

|  |  | Gai:β-arrestin-2 | CXCR3:β-arrestin-2 | CXCR3:Gai | cAMP Inhibition |
| --- | --- | --- | --- | --- | --- |
| CXCL9 | logEC50 | N/A | -7.7 ± 0.40 | -7.6 ± 0.57 | -7.9 ± 0.19 |
|  | E<sub>max</sub> | N/A | 35 ± 9.0 | 79 ± 30 | 87 ± 10 |
| CXCL10 | logEC50 | N/A | -8.5 ± 0.17 | -9.1 ± 0.55 | -10 ± 0.15 |
|  | E<sub>max</sub> | N/A | 76 ± 6.0 | 76 ± 19 | 110 ± 7.3 |
| CXCL11 | logEC50 | -7.6 ± 0.29 | -8.2 ± 0.16 | -8.8 ± 0.44 | -9.6 ± 0.21 |
|  | E<sub>max</sub> | 100 ± 20 | 100 ± 8.8 | 100 ± 21 | 100 ± 8.9 |
| VUF10661 | logEC50 | -6.0 ± 0.18 | -6.4 ± 0.17 | -6.3 ± 0.21 | -7.2 ± 0.26 |
|  | E<sub>max</sub> | 94 ± 10 | 92 ± 7.8 | 230 ± 25 | 160 ± 19 |
| VUF11418 | logEC50 | -5.9 ± 0.25 | -6.7 ± 0.28 | -5.7 ± 0.43 | -6.6 ± 0.23 |
|  | E<sub>max</sub> | 85 ± 13 | 52 ± 6.8 | 160 ± 43 | 160 ± 18 |

**Table 2:**
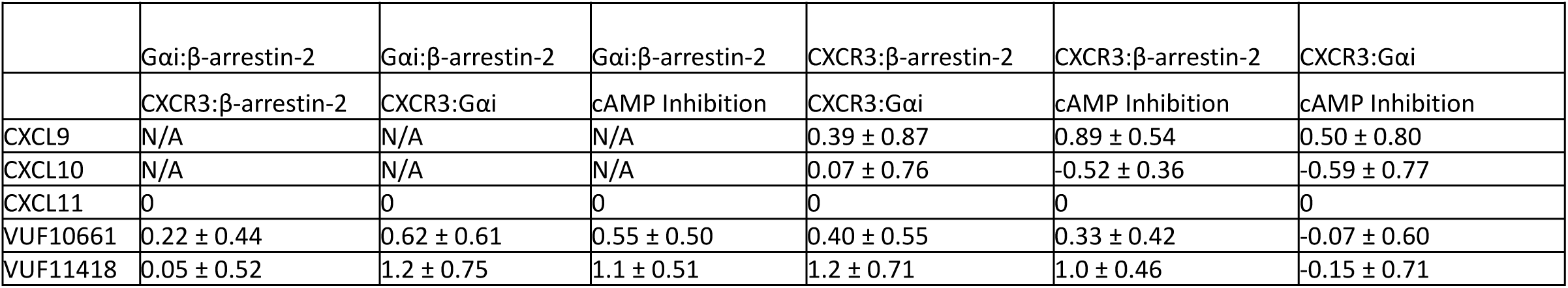
Quantified bias factors comparing signaling through different pathways. Biased factors are calculated via the logarithm of ratios of relative intrinsic activities method (see Methods).

### Biased agonists of CXCR3 induce differential conformational signatures in β-arrestin

To better characterize the effects of CXCR3 activation by biased agonists on the β-arrestin-2 structure, we used a panel of previously described FlAsH BRET conformational biosensors of β-arrestin-2 (*30*). Each FlAsH BRET biosensor features full-length *Renilla luciferase* (RLuc) at the N terminus of the β-arrestin, as well as a six-amino-acid motif, CCPGCC, at different specified residues (Fig. 4a,b). Each biosensor is able to perform intramolecular BRET between the *RLuc* and FlAsH acceptor to report on conformations adopted by β-arrestin-2 upon agonist stimulation. We modeled the positions of each FlAsH acceptor in the structures of inactive (*31*) and active (*32*) β-arrestin-1, with colored tags representing each FlAsH probe location (Fig 4b,c).

**Figure 4:**
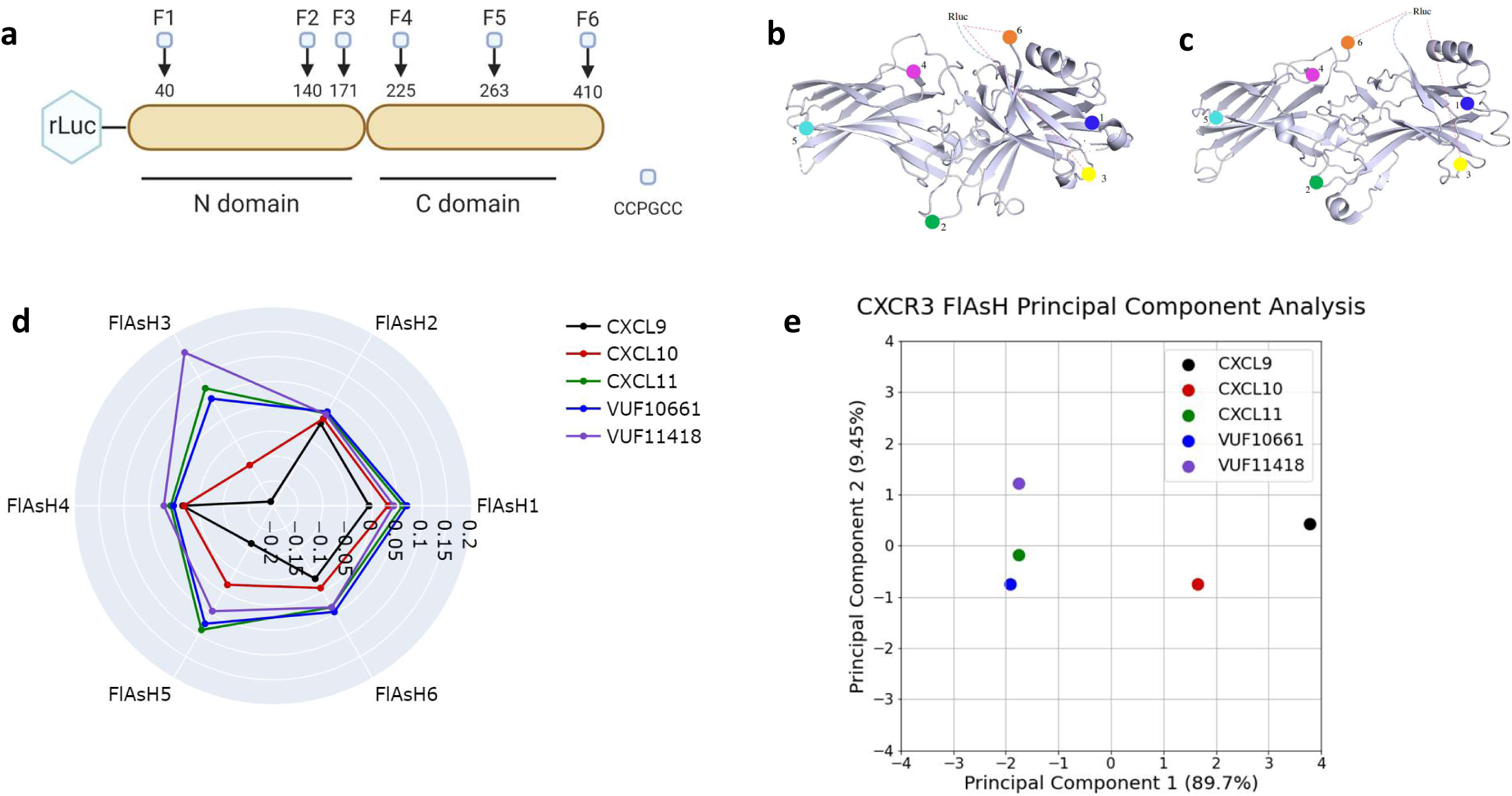
Biased agonists that produce G_αi_:β-arrestin association induce common β-arrestin conformations. A) RLuc–β-arrestin-2-FlAsH1-6 reporters (F1-F6) have amino acid motif CCPGCC inserted after β-arrestin-2 residues 40, 140, 171, 225, 263, and 410, which act as dipole acceptors to assess conformational changes in β-arrestin-2. Positions of the FlAsH binding motifs are highlighted in B) inactive and C) active structures of β-arrestin. D) HEK 293N cells were transfected with CXCR3 and either RLuc–β-arrestin2-FlAsH1, FlAsH2, FlAsH3, FlAsH4, FlAsH5, or FlAsH6. Cells then received treatment with either CXCL9 (100nM), CXCL10 (100nM), CXCL11 (100nM), VUF10661 (1μM), VUF11418 (1μM), or vehicle. Radar plot depicts intramolecular net BRET ratio calculated from subtracting vehicle from treatment group. E) Principal Component Analysis (PCA) was used to assess biased agonist induced β-arrestin conformation similarity; computed principal components are visualized with the top two principal components. Principal Component 1 contributes to 89.7% of observed variation and Principal Component 2 contributes to 9.45% of observed variation. Points denote the composite response of a single ligand at varying doses. n = 18 technical/biological replicates, graphs show mean.

In all FlAsH probes other than FlAsH2, significant differences in net BRET were observed between ligands (Fig. 4d). In both FlAsH3 and FlAsH6, significant differences were observed only between CXCL9 and CXCL10 versus CXCL11, VUF10661, and VUF11418. (Supp. Fig. 6c,f). In all FlAsH probes, no significant differences were observed between CXCL9 and CXCL10 or between CXCL11, VUF10661, and VUF11418 (Supp. Fig. 6a-f). The overall signal that we observed is more consistent with an active β-arrestin-1 structure based on the change in FlAsH3 and FlAsH 5 signal, although the FlAsH6 signal is more difficult to interpret due to the significant flexibility of the N- and C-termini of β-arrestin. Moreover, the conformation that is being induced by these ligands may be affected by the presence of G_αi_ and G_αi_:β-arrestin complex formation.

We then performed a PCA, with intramolecular BRET of each ligand for all FlAsH probes as input components (Fig. 4e). PCA captured 99% of total observed variance of the observed β-arrestin-2 conformational signatures. The conformational changes of β-arrestin-2 induced by CXCL11, VUF10661, and VUF11418 are extremely similar, with nearly identical PC1 (captures 89.7% of total variance) values. In contrast, both CXCL9 and CXCL10 are grouped much further from the other three ligands. In short, ligands that promoted G_αi_:β-arrestin complex formation also generated similar β-arrestin conformations that shared features with activated β-arrestin.

### CXCL11, VUF10661, and VUF11418 all catalyze the formation of Gαi:β-arrestin:CXCR3 complexes

We previously demonstrated that G_αi_:β-arrestin scaffolds could form selective complexes with the canonically G_αs_-coupled V_2_R and β_2_AR following stimulation with an endogenous agonist (Smith and Pack et al., submitted). However, it is unknown if canonically G_αi_-coupled receptors can form scaffolds with G_αi_:β-arrestin complexes. To address this, we utilized the three CXCR3 agonists that were observed to form the G_αi_:β-arrestin complex, CXCL11, VUF10661, and VUF11418, in a ‘complex BRET’ assay. Complex BRET is similar to other BRET-based strategies to assess complex formation (*33, 34*) and requires complementation of a low affinity nanoBiT split luciferase system followed by energy transfer to a third protein tagged with a fluorescent protein acceptor, monomeric Kusabira Orange (mKO), generating a BRET response (Fig. 5a). Thus, this technique enables real-time assessment of interactions between a two-protein complex and a third protein in living cells. Treatment with all three ligands forming G_αi_:β-arrestin complexes (CXCL11, VUF10661, and VUF11418) catalyzed the formation of G_αi_:β-arrestin:CXCR3 tripartite complexes (Fig. 5b). Although VUF10661 and CXCL11 produced the strongest G_αi_:β-arrestin signal, the rank order of G_αi_:β-arrestin:CXCR3 E_max_ changed such that VUF10661 and VUF11418 produced a greater net BRET ratio than CXCL11. As complex BRET signal depends both on distance and upon orientation, these results could indicate that these ligands are generating distinct conformations of G_αi_:β-arrestin complexes with CXCR3.

**Figure 5:**
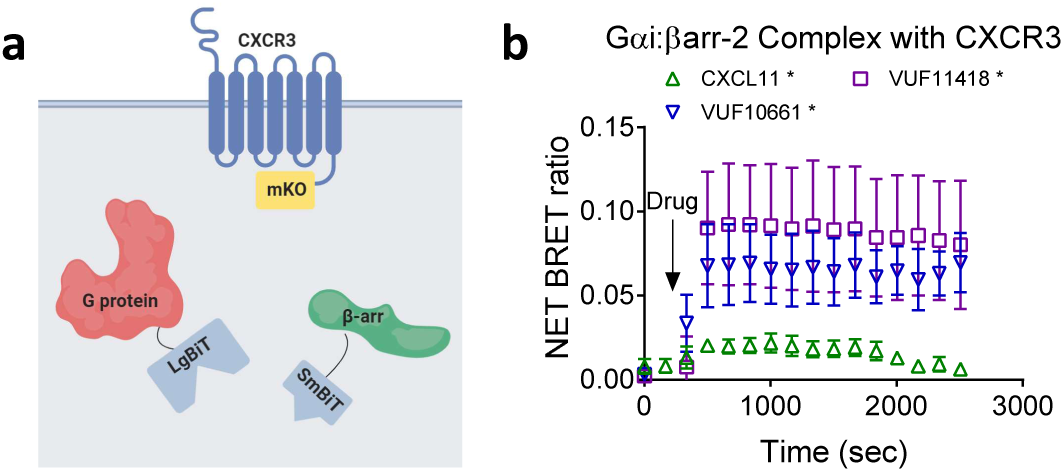
Formation of G_αi_:β-arrestin:CXCR3 tripartite complex. A) Arrangement of luciferase fragments and mKO acceptor fluorophore for complex BRET on G_αi_ (LgBiT), β-arrestin-2 (SmBiT), and CXCR3 (mKO). HEK293T cells transiently transfected with CXCR3-mKO, G_αi_-LgBiT, and SmBiT-β-arrestin-2 and stimulated with either CXCL11 (100nM), VUF10661 (1μM), VUF11418 (1μM), or vehicle. B) Complex BRET ratio for G_αi_:β-arrestin:CXCR3 tripartite complex following CXCL11, VUF10661, or VUF11418 treatment. *P<0.05 by two-way ANOVA, Dunnett’s post hoc analysis with a significant difference between treatments. n = 3-5, graphs show mean ± s.e.m.

### CXCR3 does not promote β-arrestin-dependent ERK phosphorylation or the formation of G_αi_:β-arrestin:ERK complexes

Different GPCRs regulate ERK through different mechanisms involving G proteins and β-arrestins (*6, 35, 36*). Our prior work has shown agonist treatment of the V_2_R catalyzes formation of G_αi_:β-arrestin:ERK complexes that were associated with ERK phosphorylation (Smith and Pack et al., submitted). CXCR3 activation also increases ERK1/2 phosphorylation, with CXCL9, CXCL10, and CXCL11 known to activate ERK1/2 (*21, 37*). Given that CXCL9, CXCL10, and CXCL11 can all activate ERK, but only CXCL11 was observed to induce formation of G_αi_:β-arrestin complexes, this suggested that G_αi_:β-arrestin complex formation is not necessary for activation of ERK1/2 downstream of CXCR3.

We proceeded to test the ability of CXCL11, as well as VUF10661, to form G_αi_:β-arrestin:ERK complexes (Fig. 6a). Unlike the V_2_R, agonist treatment of CXCR3 did not induce formation of G_αi_:β-arrestin:ERK complexes (Fig. 6b). We then investigated the requirement of β-arrestin-2 for CXCL11-induced activation of ERK. Unlike the V_2_R, where either CRISPR/Cas9 knockout of β-arrestin or siRNA knockdown significantly reduced ERK phosphorylation (*6*), knockout or knockdown of β-arrestin did not reduce CXCR3-dependent ERK phosphorylation (Fig 5c-j). Rather, β-arrestin knockdown or knockout increased ERK phosphorylation. Additionally, we confirmed β-arrestin siRNA knockdown, β-arrestin CRISPR/Cas9 knockout, and CXCR3 surface expression on tested cell lines (Supp. Fig. 7a-e). Taken together, these findings further support findings that ERK activation is a complex process regulated by both G proteins and β-arrestins, but does not universally depend on G_αi_:β-arrestin:ERK complex formation.

**Figure 6:**
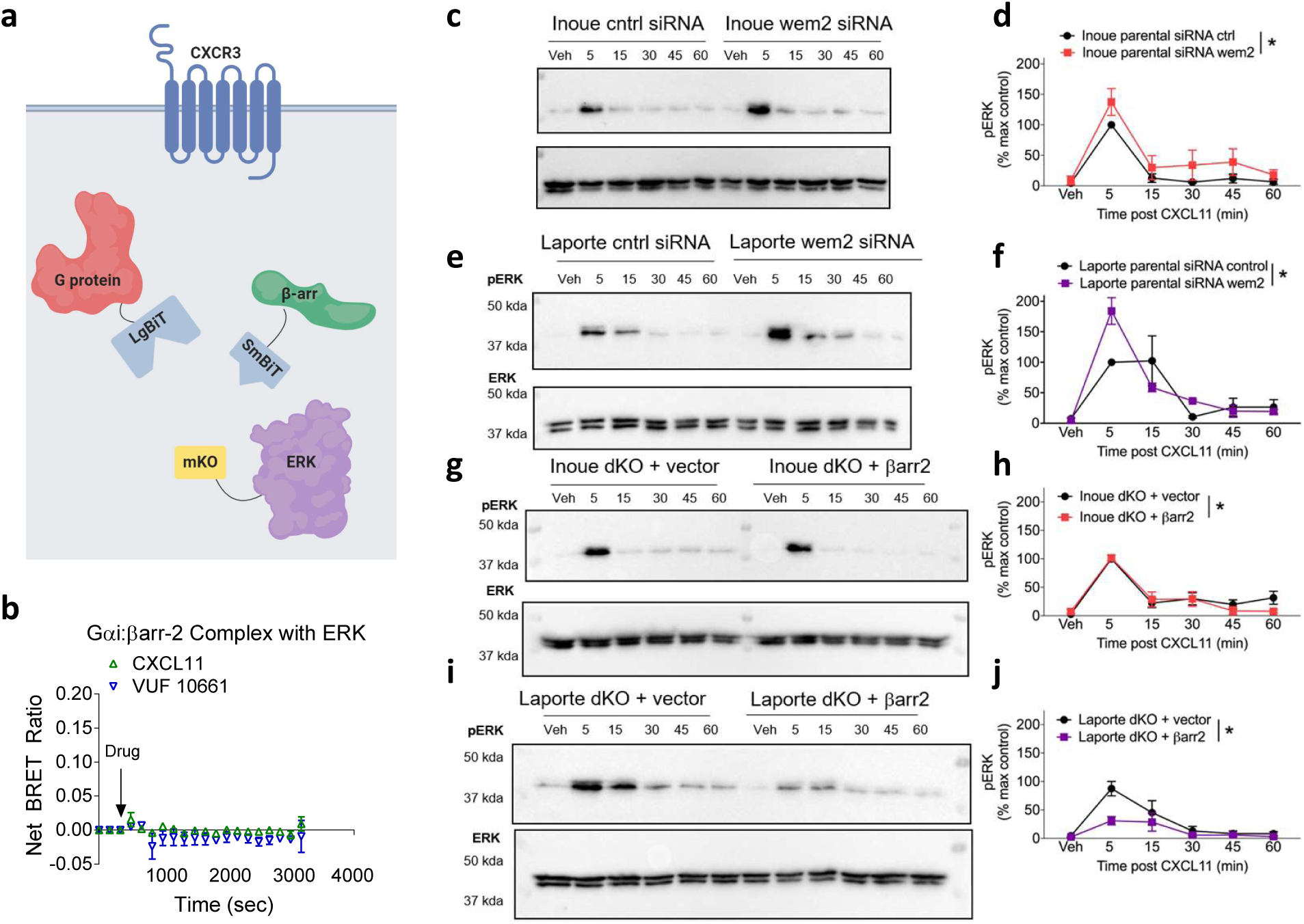
β-arrestin-2 is not necessary for CXCR3-regulated ERK activation in two cell-lines and no Gαi:β-arrestin-2:ERK complex is observed. A) Arrangement of luciferase fragments and mKO acceptor fluorophore for complex BRET on G_αi_ (LgBiT), β-arrestin (SmBiT), and ERK2 (mKO). B) Complex BRET ratio for G_αi_:β-arrestin-2:ERK following treatment with either CXCL11 (100nM), VUF10661 (1μM), or vehicle. Data were normalized to both vehicle treatment and cytosolic mKO. No complex formation between G_αi_:β-arrestin and ERK was observed. AI cells lacking β-arrestin either with C) siRNA or G) CRISPR/Cas9 knockdown of β-arrestin-1/2 resulted in increased ERK phosphorylation following five-minute treatment with CXCL11 (100 nM). Rescue of β-arrestin-2 in cells lacking β-arrestin1/2 resulted in no effect in ERK phosphorylation. Quantified pERK/ERK response in D) AI parental cells treated with control siRNA or siRNA targeted to β-arrestin-2 and H) AI β-arrestin-1/2 CRISPR/Cas9 double knockout cells (dKO). SL cells lacking β-arrestin either with E) siRNA or I) CRISPR/Cas9 knockdown of β-arrestin-2 resulted in increased ERK phosphorylation following five-minute treatment with CXCL11. Rescue of β-arrestin-2 in cells lacking β-arrestin1/2 resulted in a decrease in ERK phosphorylation. Quantified pERK/ERK response in F) SL parental cells treated with control siRNA or siRNA targeted to β-arrestin-2 and J) SL β-arrestin-1/2 CRISPR/Cas9 dKO cells. *P<0.05, two-way ANOVA, main effect of either siRNA or rescue. n = 3 replicates per condition.

## Discussion

Our results demonstrate that biased agonists of CXCR3 differentially promote the formation of G_αi_:β-arrestin complexes. We also show that CXCR3 agonists have discernable pluridimensional efficacy through canonical signaling pathways as well as G_αi_:β-arrestin complex formation. All five CXCR3 agonists inhibited cAMP and recruited G_αi_ and β-arrestin, albeit to different degrees. However, only three of the five agonists tested (CXCL11, VUF10661, and VUF11418) catalyzed G_αi_:β-arrestin complex formation, with CXCL9 and CXCL10 displaying no ability to form these complexes. Our findings provide additional evidence that G_αi_:β-arrestin complex formation is a distinct GPCR signaling pathway separable from canonical GPCR activation events.

Utilizing bias plots, along with traditional E_max_ and EC_50_ calculations, we show that these five CXCR3 agonists have distinct signaling profiles, and that activity in one signaling pathway does not necessarily predict activity in another pathway. However, we do observe some pathways with a high degree of correlation, for example, a strong degree of positive correlation between cAMP inhibition and G protein recruitment, most easily visualized within the bias plots of Figure 3. When comparing cAMP inhibition to G protein recruitment, we observe (at non-saturating concentrations of agonist) more cAMP inhibition relative to G protein recruitment. This is likely due to differences in amplification between these assays, consistent with decades of work demonstrating hormone amplification, e.g. GPCRs catalyze guanine exchange at G proteins, which can in turn catalyze second messenger signaling such as cAMP (*38, 39*).

In comparing cAMP inhibition to β-arrestin recruitment, CXCL11 and CXCL10 demonstrate bias towards β-arrestin recruitment relative to the other ligands (i.e., increased β-arrestin recruitment correlates with decreased cAMP inhibition). Similarly, the two synthetic compounds, VUF10661 and VUF11418 demonstrated greater cAMP inhibition with reduced β-arrestin recruitment. In our analyses, VUF11418 appeared to be G protein biased relative to VUF10661, consistent with prior findings using different methods (*24*), but with additional nuances of additional signaling pathway potency and efficacy now appreciated. Further analyses showed that CXCL11 is G_αi_:β-arrestin biased relative to cAMP inhibition compared to the two small molecules. As might be expected with a positive correlation between G_αi_ recruitment and cAMP inhibition, CXCL11 is also biased towards G_αi_:β-arrestin complex formation relative to G protein recruitment. In comparison to VUF10661, CXCL11 generated equal amounts of G_αi_:β-arrestin association, but much less G protein recruitment. To reduce the complexity of these signaling profiles for each ligand, we dissected these results via PCA, allowing us to demonstrate that the signaling of CXCL9 and CXCL10 were similar to one another, while CXCL11 and VUF11418 were also similar to each other. VUF10661’s signaling characteristics were extremely different from that of the two pairings. Notably, VUF10661 and VUF11418 are known to differentially regulate chemotaxis and inflammation, consistent with these signaling pathways being physiologically relevant (*24*). Overall, these findings illuminate signaling granularity within the activity of CXCR3 agonists. Our findings suggest that detailed signaling analyses of multiple pathways is likely necessary to design a drug capable of targeting CXCR3 with desired efficacy.

Using a panel of established β-arrestin biosensors (*30*), we correlate signaling activity with structural changes in β-arrestin. Consistent with bias calculations and pathway activity, CXCL9 and CXCL10 displayed different β-arrestin conformational patterns relative to CXCL11, VUF10661, and VUF11418. These data further support distinctive conformational signatures adopted by β-arrestin when CXCR3 is stimulated by biased agonists, consistent with prior reports at other GPCRs (*30, 40-42*). These agonist-induced β-arrestin conformational changes suggest that the conformational changes reported on by the FlAsH3 and FlAsH6 biosensors may be the necessary β-arrestin conformations needed to form G_αi_:β-arrestin complexes. Conformational differences in β-arrestin may thus explain in-part the inability for CXCL9 or CXCL10 to form G_αi_:β-arrestin complexes, although delineating finer structural details is necessary to test this hypothesis. Such conformational changes in β-arrestin likely exist in the absence of receptor as recently described (*43, 44*), and may also correlate with spatially distinct pools of GPCR signaling (*45, 46*).

Pertussis toxin inactivates canonical G_αi_ signaling, allowing for dissection of certain G protein and β-arrestin contributions to signaling events. While pertussis toxin had no effect on CXCR3-mediated β-arrestin recruitment, pertussis toxin reduced G_αi_ recruitment and altered G_αi_:β-arrestin association. This suggests that both G protein recruitment and G_αi_:β-arrestin association may be partially governed by underlying mechanisms disrupted by pertussis toxin. This could include a disruption of a critical interaction between alpha-5-helix of G_αi_ with CXCR3. G protein alpha-5-helix interactions with the receptor core are thought to be conserved across GPCRs and critical for disrupting key G protein:GDP contacts and initiating canonical G protein signaling (*47-49*). However, attenuation of signal and lack of elimination of G_αi_:β-arrestin complex formation with pertussis toxin pretreatment suggests an alternative interaction site beyond the alpha-5-helix, as pertussis toxin might be altering the conformational state of G_αi_.

Similar to our prior work with the G_αs_-coupled V_2_R and β_2_AR (Smith and Pack et al., submitted), G_αi_:β-arrestin complexes formed tripartite complexes with CXCR3. This is the first time to our knowledge that a G_αi_-coupled receptor has been observed to form measurable G protein:β-arrestin:receptor complexes. While all agonists capable of forming G_αi_:β-arrestin complexes also formed G_αi_:β-arrestin:CXCR3 complexes, the E_max_ rank order of ligands differed. CXCL11 was unable to form receptor tripartite complexes to the same degree as VUF11418 and VUF10661, despite producing demonstrably greater G_αi_:β-arrestin association than VUF11418. These data suggest conformational differences in how CXCL11, VUF10661, and VUF11418 induce G_αi_:β-arrestin:CXCR3 complexes, with VUF10661 and VUF11418 producing a more similar tripartite complex conformation than CXCL11. It is likely that these agonists induce different CXCR3 free energy landscapes, similar to that observed at other GPCRs (*50-53*), but further work will be necessary to confirm this.

Unlike our prior work where we showed that G_αi_:β-arrestin complexes with ERK were formed downstream of the V_2_R (Smith and Pack et al., submitted), CXCR3 did not demonstrate a similar pattern. No G_αi_:β-arrestin:ERK tripartite complexes were observed with any of the ligands we tested. While not similar to the V_2_R, it is consistent with previously described CXCR3 signaling, where CXCL9, CXCL10, and CXCL11 have been shown to activate ERK1/2 (*21, 37*). Given that only CXCL11 treatment of CXCR3 results in formation of G_αi_:β-arrestin complexes, it is likely that CXCR3 activation signals through ERK via a different mechanism than the V_2_R. Importantly, our findings demonstrate that ERK phosphorylation does not universally depend upon scaffolding ERK to G_αi_:β-arrestin complexes. Further work with a panel of receptors will be necessary to establish if ligand-generated G_αi_:β-arrestin complexes with ERK are a common mechanism of ERK activation, and if established receptor properties correlate with the ability to form G_αi_:β-arrestin:ERK complexes.

In summary, our findings further demonstrate that G_αi_:β-arrestin complexes are a fundamentally distinct GPCR signaling pathway and show that biased agonists at CXCR3 have greater granularity in signaling bias than previously appreciated. We show that G_α_:β-arrestin complexes could form larger complexes with the receptor but not with ERK. Further efforts in elucidating the signaling functions of GPCRs like CXCR3 may allow for the development of future novel pharmacologic therapies.

---

**Supplemental Table 1:** R^2^ values per CXCR3 ligand for each bias plot. R^2^ values are calculated via linear regression per agonist per bias plot.

**Supplemental Figure 1: Gα proteins recruitment to CXCR3 by biased agonists with or without pertussis toxin pretreatment.** HEK 293T transiently expressing CXCR3 and either A) G_αo_ or B) G_αs_ and treated with either CXCL9 (100nM), CXCL10 (100nM), CXCL11 (100nM), VUF10661 (1μM), VUF11418 (1μM), or vehicle at the specified concentration and analyzed for 750 seconds after an initial pre-read. Effect of pertussis toxin pretreatment (200 ng/mL) on either C) G_αi_, D) G_αo_, or E) G_αs_ with either CXCL9, CXCL10, CXCL11, VUF10661, VUF11418, or vehicle at the previously specified concentrations. *P<0.05 by two-way ANOVA, Dunnett’s post hoc analysis with a significant difference between treatments. n = 3 - 4, graphs show mean ± s.e.m.

**Supplemental Figure 2: Gα proteins recruitment to CXCR3 by biased agonists with β-arrestin-2 overexpression and pertussis toxin pretreatment.** A) HEK 293T transiently expressing CXCR3-smBiT, β-arrestin-2, and A) G_αi_-LgBiT or C) G_αs_-LgBiT and treated with either CXCL9 (100nM), CXCL10 (100nM), CXCL11 (100nM), VUF10661 (1μM), VUF11418 (1μM), or vehicle at the specified concentration and analyzed for 750 seconds after an initial pre-read. Effect of pertussis toxin pretreatment (200 ng/mL) on either B) G_αi_ or D) G_αs_ with either CXCL9, CXCL10, CXCL11, VUF10661, VUF11418, or vehicle at the previously specified concentrations. *P<0.05 by two-way ANOVA, Dunnett’s post hoc analysis with a significant difference between treatments. n = 3-4, graphs show mean ± s.e.m.

**Supplemental Figure 3: Differential cAMP inhibition by CXCR3 agonists.** HEK 293T cells transiently expressing a cAMP-activated modified firefly luciferase and CXCR3 were treated with vehicle or the indicated concentrations of A) CXCL9 (100nM), B) CXCL10 (100nM), C) CXCL11 (100nM), D) VUF11418 (1μM), E) VUF10661 (1μM) or vehicle. cAMP inhibition is recorded as percent change relative to VUF10661. High concentrations of CXCR3 peptides resulted in small but measurable amounts of non-CXCR3 mediated cAMP inhibition. *P<0.05 by two-way ANOVA, Dunnett’s post hoc analysis with a significant difference between treatments.

**Supplemental Figure 4**: **Pertussis toxin does not disrupt β-arrestin recruitment to CXCR3.** HEK 293T cells transiently expressing CXCR3-LgBiT and smBiT-β-arrestin2 received pertussis toxin pretreatment (200 ng/mL) and were subsequently treated with either A) CXCL9 (100nM), B) CXCL10 (100nM), C) CXCL11 (100nM), D) VUF11418 (1μM), E) VUF10661 (1μM) or vehicle at specified concentration. No significant changes between pertussis toxin pretreated cells and non-pretreated cells were observed. n.s. by two-way ANOVA, Dunnett’s post hoc analysis with no significant difference between treatments.

**Supplemental Figure 5**: **Biased agonists of CXCR3 differentially drive formation of G**_**αo**_**:β-arrestin complexes through CXCR3, but not through G**_**αs**_, **G**_**αq**_, **or G**_**α12**_. A) HEK 293T cells transiently expressing CXCR3, β-arrestin-2, and A) G_αo_, B) G_αs_, C) G_αq_, or D) G_α12_ were treated with CXCL9 (100nM), CXCL10 (100nM), CXCL11 (100nM), VUF10661 (1μM), VUF11418 (1μM), or vehicle. Of these additional G protein families, only G_αo_ had the ability to form a G_αo_:β-arrestin complex, and significant G_αo_:β-arrestin complex formation was only observed following either CXCL11 or VUF10661 treatment. HEK293T without CXCR3 overexpression observe minor G_αi_:β-arrestin2 association when treated with CXCL9. E) Transfected HEK 293T expressing smBiT-β-arrestin2 and G_αi_-LgBit, but no CXCR3. The cells were treated with either CXCL9, CXCL10, CXCL11, VUF10661, VUF11418, or vehicle and minor recruitment was observed with only CXCL9. *P<0.05 by two-way ANOVA, Dunnett’s post hoc analysis with a significant difference between treatments. n = 3-5, graphs show mean ± s.e.m.

**Supplemental Figure 6**: **Biased agonists induce differential RLuc–β-arrestin2–FlAsH1-6 conformational signatures at CXCR3.** HEK 293N cells were transfected with CXCR3 and either RLuc–β-arrestin2-A) FlAsH1, B) FlAsH2, C) FlAsH3, D) FlAsH4, E) FlAsH5, or F) FlAsH6. Cells then received treatment with either CXCL9 (100nM), CXCL10 (100nM), CXCL11 (100nM), VUF10661 (1μM), VUF11418 (1μM), or vehicle. Graphs depict intramolecular net BRET ratio calculated from subtracting vehicle from treatment group. *P<0.05 by one-way ANOVA, Tukey-Kramer post hoc analysis. n = 18 technical/biological replicates, graphs show mean ± s.e.m.

**Supplemental Figure 7**: **siRNA knockdown and CRISPR/Cas9 knockout validation.** Knockdown of β-arrestin-2 using control or siRNA directed against β-arrestin-2 in A) AI and B) SL parental cells. β-arrestin-2 rescue in C) AI and D) SL β-arrestin-1/2 dKO cells is effective. E) AI dKO cells express significantly reduced CXCR3 at the surface relative to the Inoue parental cells. No significant difference was found in CXCR3 surface expression between the SL parental and SL dKO cells. *P<0.05, two-way ANOVA, main effect of either CRISPR/Cas9 or siRNA knockdown. n = 3 replicates per condition.

## Methods

### Cell culture and transfection

Human embryonic kidney cells (HEK 293T) were maintained in minimum essential medium supplemented with 1% anti-anti and 10% fetal bovine serum. Cells were grown at 37 °C with humidified atmosphere of 5% CO_2_. For BRET and luminescence studies, HEK 293T cells were transiently transfected via an optimized calcium phosphate protocol as previously described (*21*).

### Split luciferase and complex BRET assays

HEK293T cells seeded in 6-well plates were co-transfected with smBiT, LgBiT, and mKO tagged components as previously described (Smith and Pack et al., submitted). Twenty-four hours post-transfection, cells were plated onto clear bottom, white-walled 96-well plates at 50,000-100,000 cells/well in “BRET media” - clear minimum essential medium (GIBCO) supplemented with 2% FBS, 10 mM HEPES, 1x GlutaMax, and 1x Anti-Anti (GIBCO). Select cells were then treated overnight with pertussis toxin pretreatment at a final concentration of 200 ng/mL. For luminescence split luciferase studies, plates were read with a BioTek Synergy Neo2 plate reader set at 37 °C with a 485 nm emission filter. Cells were stimulated with either vehicle (Hank’s Balanced Salt Solution with 20 mM HEPES) or indicated concentration of agonist. For split luciferase luminescence experiments, plates were read both before and after ligand treatment to calculate Δnet change in luminescence and subsequently normalized to vehicle treatment. For complex BRET experiments, plates were read on a Berthold Mithras LB940 using pre-warmed media and instrument at 37 °C using a standard RLuc emissions filter (480 nm) with a custom mKO 542 nm long-pass emission filter (Chroma Technology Co., Bellows Falls, VT) as previously described (Smith and Pack et al., submitted).

### FlAsH Intramolecular BRET

Intramolecular BRET at β-arrestin-2 conformational biosensors was measured based on a modified version of the steps and procedures previously described (*30*). Briefly, HEK293N cells were seeded in 6-well plates at a cell density of 5×10^5^ cells/well and transfected with CXCR3 receptor and one of the six FlAsH constructs. Experiments were conducted by transfecting 2000ng of CXCR3 and 200ng of FlAsH constructs. Clear bottom, white walled 96-well plates were prepared by coating with rat collagen. Twenty-four hours post transfection, cells were plated onto these coated 96-well plates at 50,000-100,000 cells/well in minimum essential medium supplemented with 1% anti-anti and 10% fetal bovine serum. In preparation of being read, cells were treated with the biarsenical labeling reagent FlAsH-EDT2 for 45 minutes, washed with BAL Wash Buffer, and suspended in Hank’s Balanced Salt Solution with 20 mM HEPES. Cells were stimulated by either vehicle (Hank’s Balanced Salt Solution with 20 mM HEPES) or indicated concentration of agonist. Immediately before reading, cells were treated with coelenterazine and read on a Berthold Mithras LB940 using pre-warmed instrument at 37 °C using a standard RLuc emissions filter (480 nm) with a custom eYFP filter (530 nm).

### cAMP Inhibition

GloSensor cAMP inhibition was conducted similar to that previously described (*21*). GloSensor biosensor (Promega) uses a modified form of firefly luciferase containing a cAMP-binding motif. Upon cAMP binding, a conformational change leads to enzyme complementation and incubation with a luciferase substrate results in a luminescence readout. Analysis of cAMP accumulation was performed in HEK293 cells transiently transfected with the Glosensor construct and human CXCR3. Cells were seeded in 96-well white, clear-bottomed plates at 80,000 cells/well, in minimum essential medium supplemented with 1% anti-anti and 10% fetal bovine serum. The next day, the GloSensor reagent [Promega; 4% (v/v)] was incubated at room temperature for 2 h. Cells were then stimulated with a range of CXCR3 agonists for 5 min, and increases in luminescence were read on a BioTek Synergy Neo2 plate reader set at 37 °C with a 485 nm emission filter.

### Structural Protein Representations

Schematic representations for inactive (Protein Data Bank (PDB) code 1G4M) and NTSR1 bound (Protein Data Bank (PDB) code 6PWC) β-arrestin-1 were derived from inactive and active β-arrestin crystal structures. Sequence alignment of β-arrestin-1 and β-arrestin-2 was utilized to translate FlAsH reporter sequence location from β-arrestin-2 to β-arrestin-1.

### ERK Immunoblots

Parental and β-arrestin1/2 knockout HEK293 cell lines (AI and SL) were previously generated and validated (*6*). Reconstitution of β-arrestin1/2 in CRISPR/Cas9 HEK293 cells by transient transfection was conducted as previously described (*6, 21*). β-arrestin1/2 knockdown in parental HEK293 cells using siRNA was conducted using Lipofectamine 3000 (ThermoFischer) according to manufacturer specifications, similar to that previously described (*6*). AI-parental were maintained in minimal essential medium (MEM) supplemented with 10% fetal bovine serum (FBS) and 1% penicillin/streptomycin or gentamicin (20 μg/ml). SL-parental cells were maintained in Dulbecco’s modified Eagle’s medium (DMEM) without pyruvate supplemented with 10% FBS and 1% penicillin/streptomycin or gentamicin (20 μg/ml). β-arrestin-1, β-arrestin-2, phosphorylated extracellular regulated kinase 1/2 (ERK) (Cell Signaling Technology, Danvers, MA) and total ERK (Millipore, Burlington, MA) antibodies were used to assess β-arrestin-1 expression ERK activation as previously described (*21*).

### Drugs

CXCL9, CXCL10, and CXCL11 were prepared and stored according to manufacturer specifications with 0.1% bovine serum albumin utilized as a carrier protein. VUF10661 (Sigma-Aldrich) and VUF11418 (Aobious) were dissolved in dimethyl sulfoxide (DMSO) to make stock solutions and stored in a desiccator cabinet. All drug dilutions were performed with BRET media or cell culture media. Pertussis toxin was obtained from List Biological Laboratories (Campbell, CA). All compound stocks were stored at −20°C until use.

## Acknowledgements

The authors thank N. Nazo for administrative assistance; D. Eiger and C. Lee for helpful discussion and thoughtful feedback. This work was supported by T32GM7171 (J.S.S.), the Duke Medical Scientist Training Program (J.S.S.), F31DA041160 (T.F.P.), PRIME JP17gm5910013 (A.I.), the LEAP JP17gm0010004 from the Japan Agency for Medical Research and Development (A.I.), the JSPS KAKENHI (A.I.), 17K08264R37MH073853 (M.G.C.),1R01GM122798-01A1 (S.R.), K08HL114643-01A1, (S.R.), Burroughs Wellcome Career Award for Medical Scientists (S.R.).

## Author Contributions

K.Z. and J.S.S. contributed equally to this work. K.Z., J.S.S., T.F.P., M.G.C., and S.R. conceived of the study and designed experiments. A.I. designed, generated, and validated all G protein split luciferase constructs. K.Z., J.S.S., A.W., I.C., and J.N.G. performed the experiments. K.Z., J.S.S., J.N.G., and S.R. analyzed the data. K.Z., J.S.S., and S.R. wrote the paper. All authors discussed the results and commented on the manuscript.

