## Supplementals for "Biased agonists of the chemokine receptor CXCR3 differentially drive formation of G_αi_:β-arrestin complexes"

**Supp. Table 1**

| | Gαi:β-arrestin-2 | Gαi:β-arrestin-2 | Gαi:β-arrestin-2 | CXCR3:β-arrestin-2 | CXCR3:β-arrestin-2 | CXCR3:Gαi |
|---|---|---|---|---|---|---|
| | CXCR3:β-arrestin-2 | CXCR3:Gαi | cAMP Inhibition | CXCR3:Gαi | cAMP Inhibition | cAMP Inhibition |
| CXCL9 | N/A | N/A | N/A | 0.68 | 0.92 | 0.86 |
| CXCL10 | N/A | N/A | N/A | 0.87 | 0.51 | 0.55 |
| CXCL11 | 0.90 | 0.84 | 0.61 | 0.83 | 0.72 | 0.82 |
| VUF10661 | 0.95 | 0.98 | 0.72 | 0.97 | 0.83 | 0.84 |
| VUF11418 | 0.84 | 0.88 | 0.89 | 0.78 | 0.99 | 0.85 |

### Supp. 1

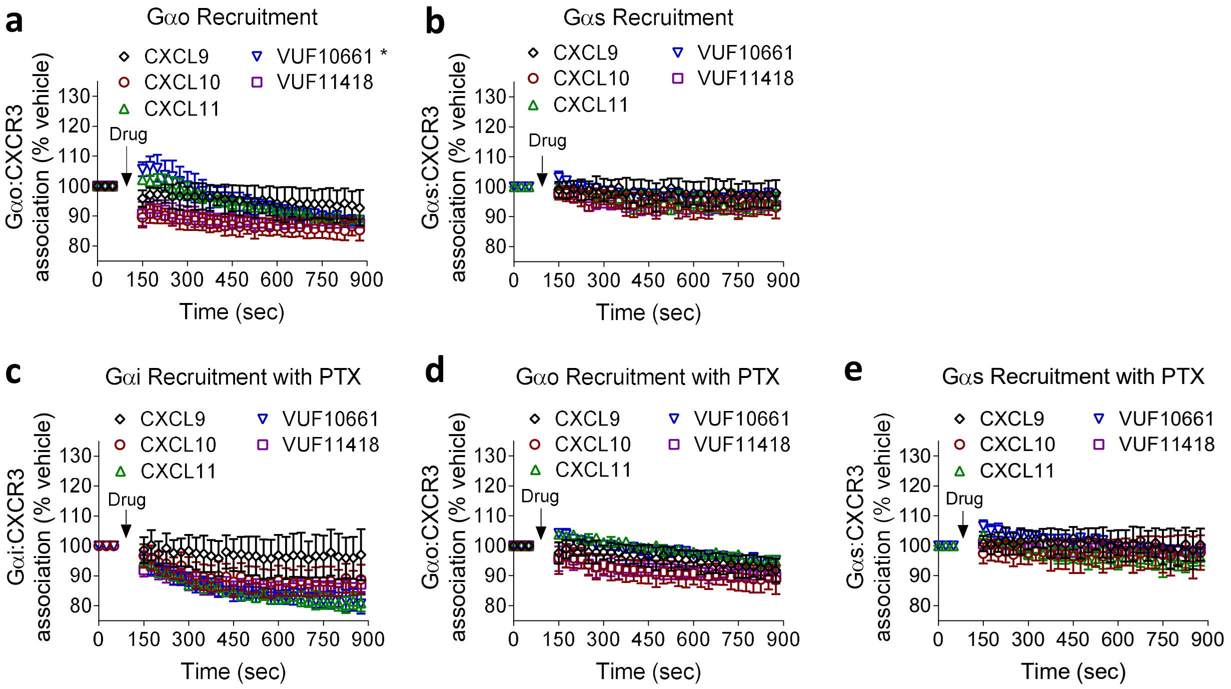
a
Gαo Recruitment
CXCL9
CXCL10
CXCL11
VUF10661 *
VUF11418
Gαo:CXCR3 association (% vehicle)
Drug
Time (sec)
b
Gαs Recruitment
CXCL9
CXCL10
CXCL11
VUF10661
VUF11418
Gαs:CXCR3 association (% vehicle)
Drug
Time (sec)
c
Gαi Recruitment with PTX
Gαi:CXCR3 association (% vehicle)
Drug
Time (sec)
d
Gαo Recruitment with PTX
Gαo:CXCR3 association (% vehicle)
Drug
Time (sec)
e
Gαs Recruitment with PTX
Gαs:CXCR3 association (% vehicle)
Drug
Time (sec)

### Supp. 2

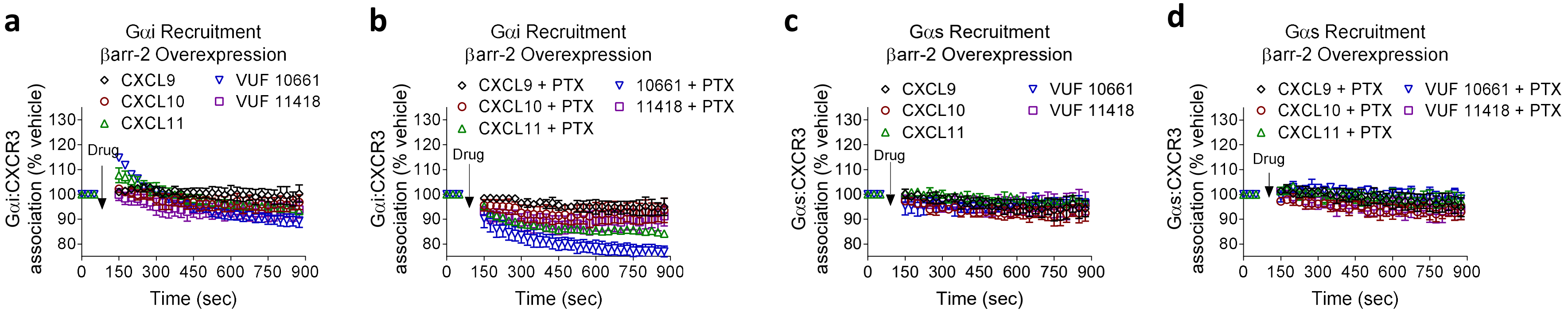

a
Gαi Recruitment
βarr-2 Overexpression
CXCL9
CXCL10
CXCL11
VUF 10661
VUF 11418
Gαi:CXCR3 association (% vehicle)
Drug
Time (sec)
b
Gαi Recruitment
βarr-2 Overexpression
CXCL9 + PTX
CXCL10 + PTX
CXCL11 + PTX
10661 + PTX
11418 + PTX
Gαi:CXCR3 association (% vehicle)
Drug
Time (sec)
c
Gαs Recruitment
βarr-2 Overexpression
CXCL9
CXCL10
CXCL11
VUF 10661
VUF 11418
Gαs:CXCR3 association (% vehicle)
Drug
Time (sec)
d
Gαs Recruitment
βarr-2 Overexpression
CXCL9 + PTX
CXCL10 + PTX
CXCL11 + PTX
VUF 10661 + PTX
VUF 11418 + PTX
Gαs:CXCR3 association (% vehicle)
Drug
Time (sec)

### Supp. 3

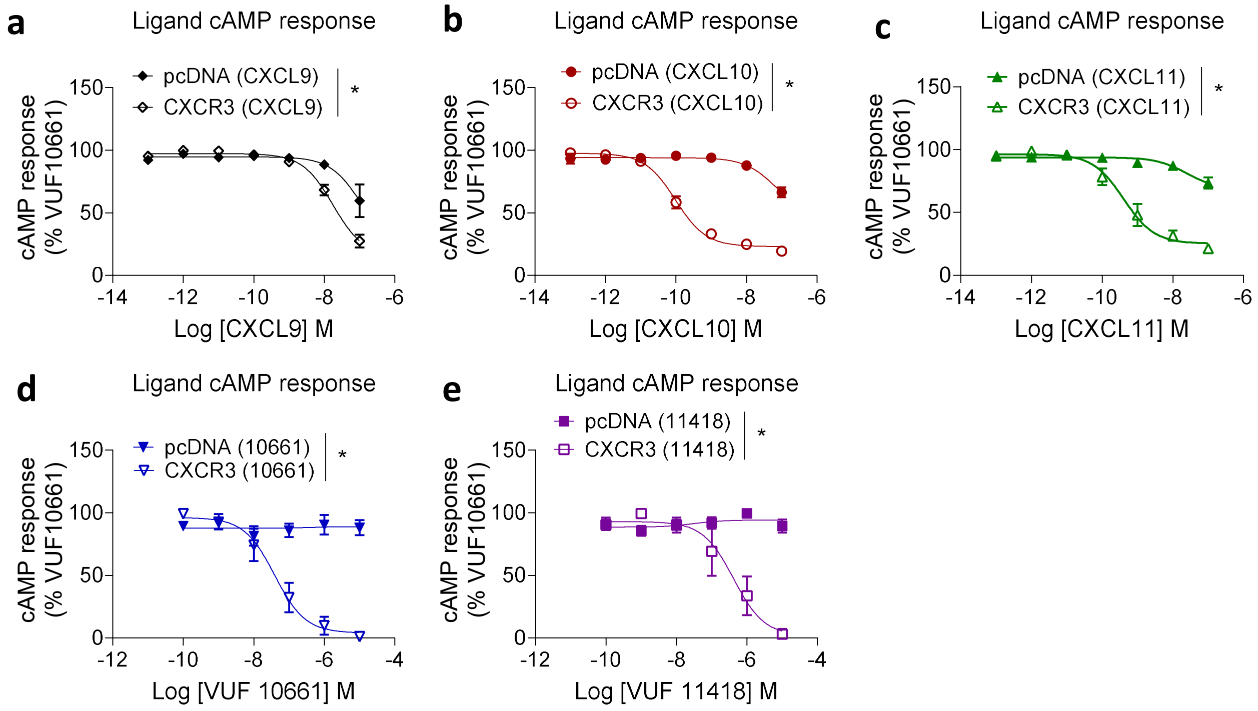
a
Ligand cAMP response
pcDNA (CXCL9)
CXCR3 (CXCL9)
*
cAMP response (% VUF10661)
Log [CXCL9] M
b
Ligand cAMP response
pcDNA (CXCL10)
CXCR3 (CXCL10)
*
cAMP response (% VUF10661)
Log [CXCL10] M
c
Ligand cAMP response
pcDNA (CXCL11)
CXCR3 (CXCL11)
*
cAMP response (% VUF10661)
Log [CXCL11] M
d
Ligand cAMP response
pcDNA (10661)
CXCR3 (10661)
*
cAMP response (% VUF10661)
Log [VUF 10661] M
e
Ligand cAMP response
pcDNA (11418)
CXCR3 (11418)
*
cAMP response (% VUF10661)
Log [VUF 11418] M

**Supp. 4**

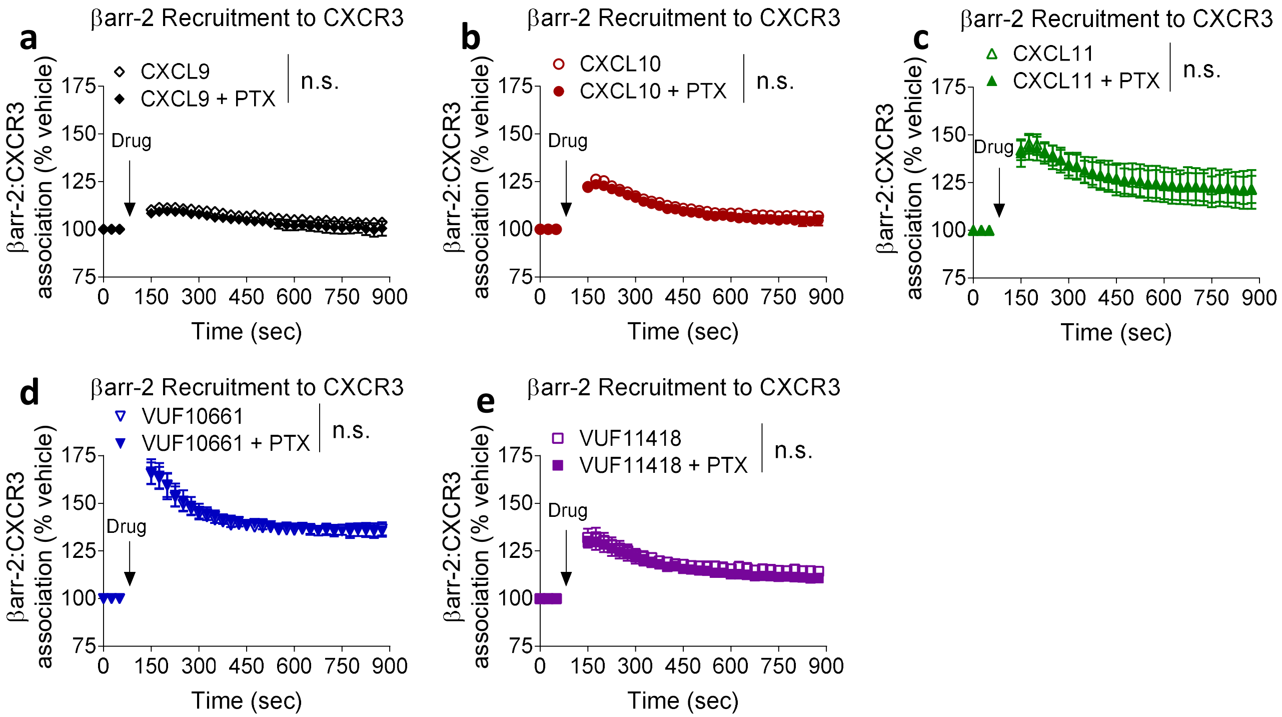

a
βarr-2 Recruitment to CXCR3
CXCL9
CXCL9 + PTX
n.s.
Drug
βarr-2:CXCR3 association (% vehicle)
Time (sec)
b
βarr-2 Recruitment to CXCR3
CXCL10
CXCL10 + PTX
n.s.
Drug
βarr-2:CXCR3 association (% vehicle)
Time (sec)
c
βarr-2 Recruitment to CXCR3
CXCL11
CXCL11 + PTX
n.s.
Drug
βarr-2:CXCR3 association (% vehicle)
Time (sec)
d
βarr-2 Recruitment to CXCR3
VUF10661
VUF10661 + PTX
n.s.
Drug
βarr-2:CXCR3 association (% vehicle)
Time (sec)
e
βarr-2 Recruitment to CXCR3
VUF11418
VUF11418 + PTX
n.s.
Drug
βarr-2:CXCR3 association (% vehicle)
Time (sec)

**Supp. 5**

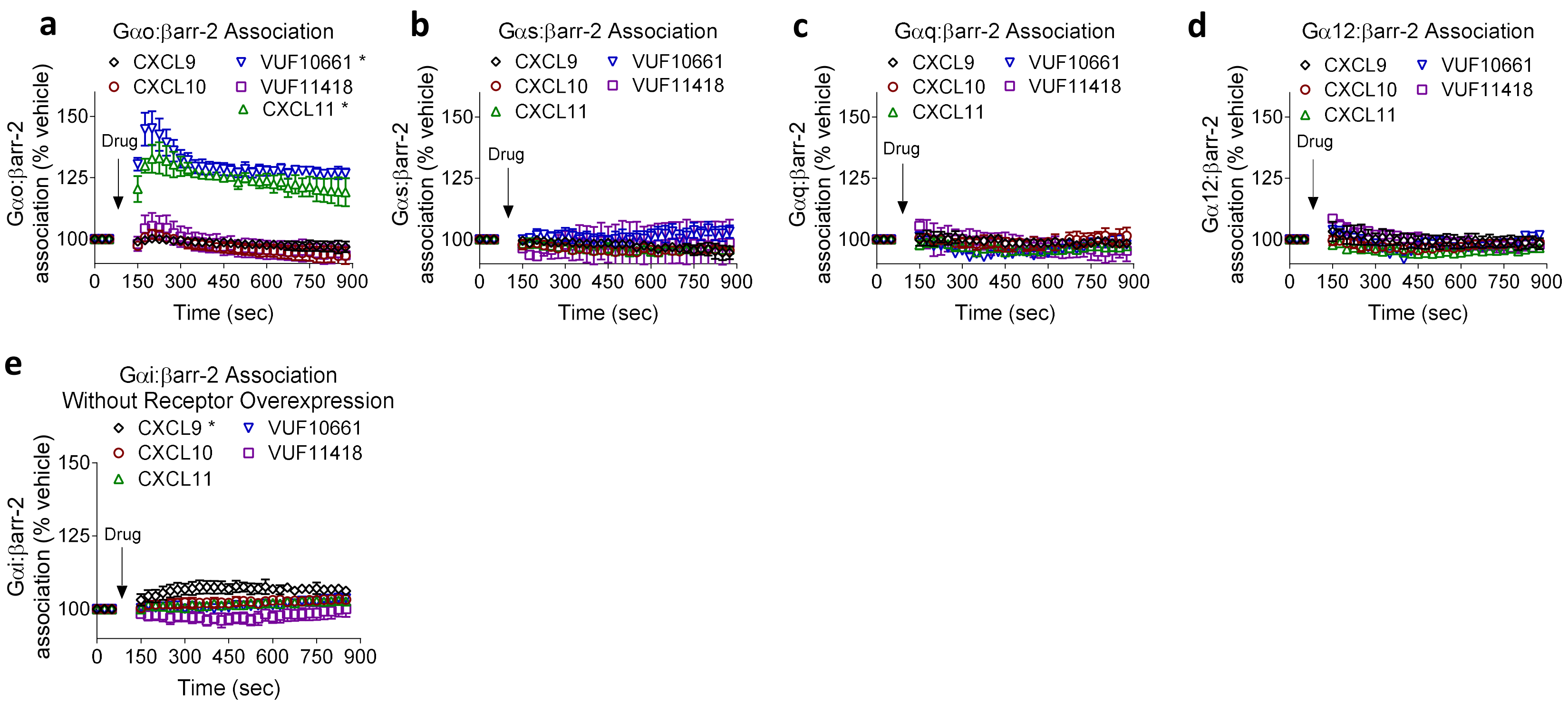

a
Gαo:βarr-2 Association
CXCL9
CXCL10
VUF10661 *
VUF11418
CXCL11 *
Gαo:βarr-2 association (% vehicle)
Drug
Time (sec)
b
Gαs:βarr-2 Association
CXCL9
CXCL10
CXCL11
VUF10661
VUF11418
Gαs:βarr-2 association (% vehicle)
Drug
Time (sec)
c
Gαq:βarr-2 Association
CXCL9
CXCL10
CXCL11
VUF10661
VUF11418
Gαq:βarr-2 association (% vehicle)
Drug
Time (sec)
d
Gα12:βarr-2 Association
CXCL9
CXCL10
CXCL11
VUF10661
VUF11418
Gα12:βarr-2 association (% vehicle)
Drug
Time (sec)
e
Gαi:βarr-2 Association
Without Receptor Overexpression
CXCL9 *
CXCL10
CXCL11
VUF10661
VUF11418
Gαi:βarr-2 association (% vehicle)
Drug
Time (sec)

Supp. 6

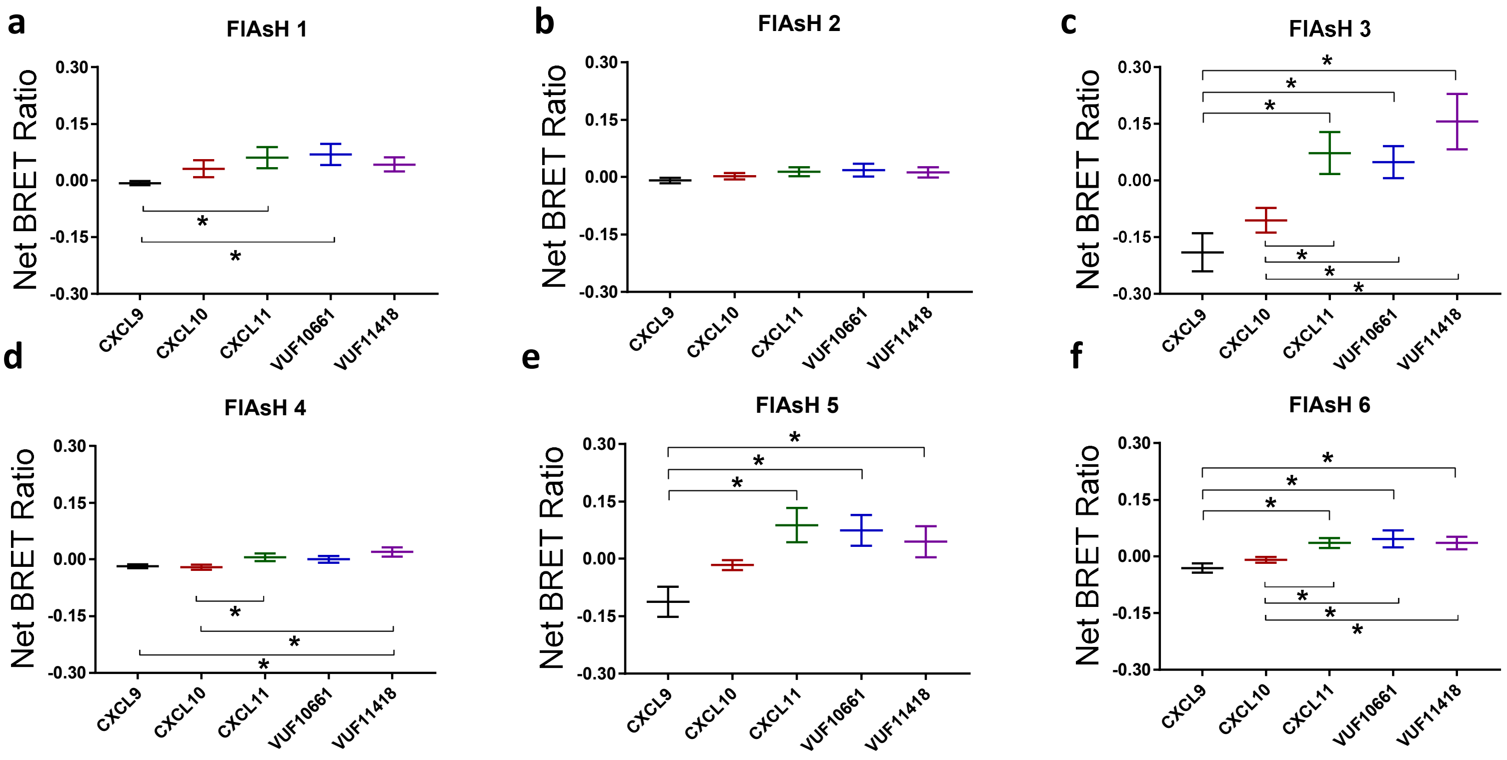

a
FlAsH 1
b
FlAsH 2
c
FlAsH 3
d
FlAsH 4
e
FlAsH 5
f
FlAsH 6
Net BRET Ratio
0.30
0.15
0.00
-0.15
-0.30
*
CXCL9
CXCL10
CXCL11
VUF10661
VUF11418

**Supp. 7**

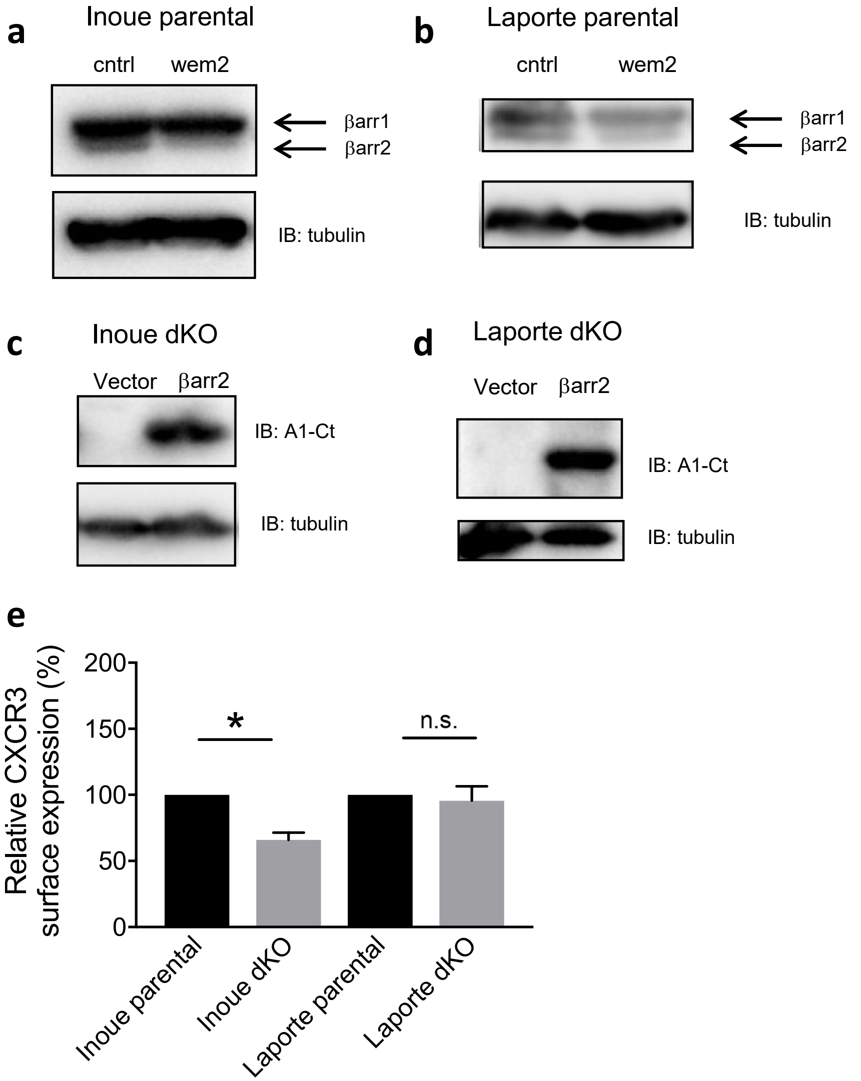

a
Inoue parental
cntrl
wem2
βarr1
βarr2
IB: tubulin
b
Laporte parental
cntrl
wem2
βarr1
βarr2
IB: tubulin
c
Inoue dKO
Vector
βarr2
IB: A1-Ct
IB: tubulin
d
Laporte dKO
Vector
βarr2
IB: A1-Ct
IB: tubulin
e
Relative CXCR3 surface expression (%)
200
150
100
50
0
*
n.s.
Inoue parental
Inoue dKO
Laporte parental
Laporte dKO
